# Provision of Preferred Nutrients to Macrophages Enables *Salmonella* to Replicate Intracellularly Without Relying on Type III Secretion Systems

**DOI:** 10.1101/2025.05.15.653970

**Authors:** Francisco-Javier Garcia-Rodriguez, Camila Valenzuela, Joaquin Bernal-Bayard, Pedro Escoll

## Abstract

Intracellular survival and replication within macrophages are key virulence determinants of *Salmonella enterica* serovar Typhimurium. This phenomenon is traditionally attributed to the activity of its two Type III Secretion Systems (T3SS) and their associated effectors. A critical challenge for these bacteria is acquiring nutrients from inside the host cell. Thus, they modulate the metabolism of host cells to replicate. Given the metabolic plasticity of macrophages, a key unresolved question is how their metabolic heterogeneity shapes intracellular *Salmonella* replication. By using human primary macrophages and live-cell imaging to monitor bacterial dynamics at the single-cell level, we revealed that *Salmonella* does not replicate in all infected cells. However, supplementation with specific carbon sources used by *Salmonella* during infection accelerated bacterial replication and increased the proportion of macrophages showing replicative bacteria. Remarkably, this occurred even in the absence of functional T3SSs, as a Δ*prgH*/Δ*ssaV* double mutant was able to replicate in a subset of infected cells under favorable nutrient conditions. These phenotypes are further amplified in macrophages with higher glycolytic activity, such as the murine RAW 264.7 cell line. Further analyses demonstrated that enhanced *Salmonella* replication is not strictly dependent on host glycolytic activity but is instead driven by the ability of the host cell to take up the nutrients *Salmonella* prefers for its replication early during infection. In summary, our findings suggest that the dependence of *Salmonella* on its T3SSs for intracellular replication can be bypassed when host cells provide optimal access to key nutrients and highlight the impact of metabolic heterogeneity in shaping intracellular bacterial replication during infection of macrophages.

## Introduction

Non-typhoidal *Salmonella* (NTS) are foodborne pathogens and a leading cause of gastroenteritis in humans and animals, affecting approximately 93.8 million people and causing 155,000 fatalities globally each year. ^1^ While most cases in healthy humans consist of a self-limiting gastroenteritis, it can also cause systemic infection in immunocompromised humans as well as very young or older individuals. Systemic infections with NTS are difficult to treat and are associated with a 15% case-fatality ratio. ^2^ *Salmonella enterica* serovar Typhimurium (ST) is one of the most common NTS serovars isolated from the bloodstream of patients. ^3,4^ In the systemic disease caused by ST, the ability to survive and replicate in macrophages is essential to the systemic spread of the bacteria. Within macrophages, ST reaches organs via the lymphatic system, such as the liver, the gallbladder, or the spleen, and access the bloodstream. ^5–7^

Virulence of ST relies on the expression of two secretion systems encoded by the *Salmonella* pathogenicity islands (SPI)-1 and SPI-2. The SPI-1 encodes a type III secretion system (T3SS1) that secretes bacterial effectors facilitating bacterial entry into non-phagocytic cells. ^8,9^ The SPI-2 encodes the T3SS2, which secretes a group of bacterial effectors, once ST invades eukaryotic cells, to generate and maintain a modified phagosome named the *Salmonella*-containing vacuole (SCV) where ST survives and replicates during infection. The T3SS2 and several secreted effectors are required for the survival and replication of ST within the SCV in murine and human primary macrophages, as well as in macrophage cell lines, and is also required for *Salmonella* virulence in mice. ^5,10–12^

Access to host nutrients in infected tissues is fundamental for *Salmonella* growth and virulence. Steeb and colleagues demonstrated that ST does not rely on a single dominant nutrient but rather exploits multiple host-derived carbon sources in parallel to sustain its intracellular replication and systemic infection. They identified glycerol, glucose, and lactate, among others, as key nutrients that contribute to *Salmonella* host tissue colonization. ^13^ Other studies showed that ST preferentially utilizes glucose as its main carbon source during intracellular replication, particularly within the SCV formed in macrophages and epithelial cells. ^14^ ST mutants defective in glycolysis and glucose uptake exhibit reduced replication rates in macrophage cell lines, highlighting the necessity of glucose metabolism for efficient intracellular growth. However, these mutants can still replicate, albeit less efficiently, suggesting that ST can switch to other carbon sources in the absence of glucose, possibly utilizing C3 substrates such as glycerol. ^14,15^ In systemically infected mice, where ST replicates mainly in macrophages, bacterial growth depends on a complete tricarboxylic acid (TCA) cycle of ST, where mutants unable to convert succinate to oxaloacetate are avirulent, highlighting the critical role of central carbon metabolism in virulence. ^14,16^ Moreover, ST adapts bacterial metabolism to the type of macrophage it infects.

In M2-polarized macrophages, which are anti-inflammatory and mainly use lipid oxidation, ST exploits increased glucose availability to support its growth. ^17,18^ In contrast, in M1-polarized macrophages, which are pro-inflammatory and rely on glycolysis, ST switches its metabolism to oxidize fatty acids, using lipid transport systems to maintain survival. ^17,19,20^ Recent findings further highlight that ST induces macrophage polarization and metabolic reprogramming, contributing to an intracellular environment compatible with bacterial survival and replication. ^21,22^

To further understand the impact of macrophage metabolism in the intracellular replication of ST, we restricted human primary macrophages to different bioenergetic contexts by culturing them in media containing specific carbon nutrients sources, as extracellular metabolites can reach intracellular *Salmonella* and contribute to their nutrition. ^13,23,24^ We analyzed macrophage metabolism during ST infection and followed bacterial replication rate at the single cell level. Our results revealed that ST can still replicate intracellularly in the absence of any T3SS and independently of host cell glycolysis when specific carbon sources are available in the cytoplasm of the infected cell. This suggests that the two T3SSs and their bacterial effectors are not required for the intracellular replication of ST if the bacterium finds a favorable metabolic environment within macrophages.

## Results

### Glycerol enhances intracellular growth dynamics of *Salmonella* Typhimurium within human primary macrophages

To study the impact of the metabolic heterogeneity of macrophages on ST intracellular replication we need to capture infection dynamics at the single-cell level. We infected human monocyte-derived macrophages (hMDMs) with GFP-expressing ST wild-type (ST-WT) and acquired time-lapse images of thousands of living macrophages at hourly intervals up to 16 hours postinfection (hpi) **(Figure 1A)**. Using our image analysis pipeline (see Methods section and ref. ^25^), we tracked individual infected cells and quantified bacterial replication by measuring the area occupied by intracellular ST-WT across time-lapse experiments **(Figure 1B)**. As ST virulence depends on the function of both Type-III Secretion Systems (T3SSs), we infected hMDMs with an isogenic ST-Δ*prgH*/Δ*ssaV* double mutant, which is T3SS-deficient and served as control for bacterial replication measurements **(Figure 1C)**. This approach allowed us to distinguish between macrophages supporting ST growth and those where ST did not replicate. To assess the role of specific nutrients in supporting ST growth within macrophages, we followed bacterial dynamics in hMDMS cultured in nutrient-limited medium supplemented with key metabolites that ST exploits for intracellular growth during *in vivo* infection, including glucose, glycerol, lactate, glutamine, and pyruvate. ^13–15^ As control, hMDMs were maintained in the same nutrient-limited medium without any additional carbon source. Our single-cell analyses of bacterial dynamics in tracked, infected cells showed that, when hMDMs were cultured in glycerol, they exhibited a higher rate at which the area occupied by intracellular bacteria increased over time compared to glucose, lactate, glutamine, pyruvate and control conditions **(Figure 1D** and **S1C)**. Moreover, the percentage of infected macrophages with replicative ST was also significantly higher in hMDMs supplemented with glycerol than in any other culture condition (58 ± 13% in glycerol vs. 18 ± 9% in control, p<0.0001). Glucose or lactate only had a minor effect in increasing the percentage of macrophages containing replicating ST **(Figure 1E)**. These results indicated that glycerol, rather than glucose, is a preferred carbon source for ST replication within human macrophages. Surprisingly, hMDMs infected with the ST-Δ*prgH*/Δ*ssaV* mutant showed that this T3SS-deficient strain was able to replicate in approximately 4% of infected hMDMs when glycerol was present in the culture medium, but not under any other condition **(Figure 1F)**. Single cell tracking showed that glycerol increased the intracellular growth of both ST-WT and the ST-Δ*prgH*/Δ*ssaV* mutant compared to glucose conditions **(Figure 1B, 1C, 1G** and **1H)**. These findings suggested that, despite the absence of both T3SSs, ST can replicate within primary human macrophages when specific nutrients, such as glycerol, are available.

**Figure 1.**
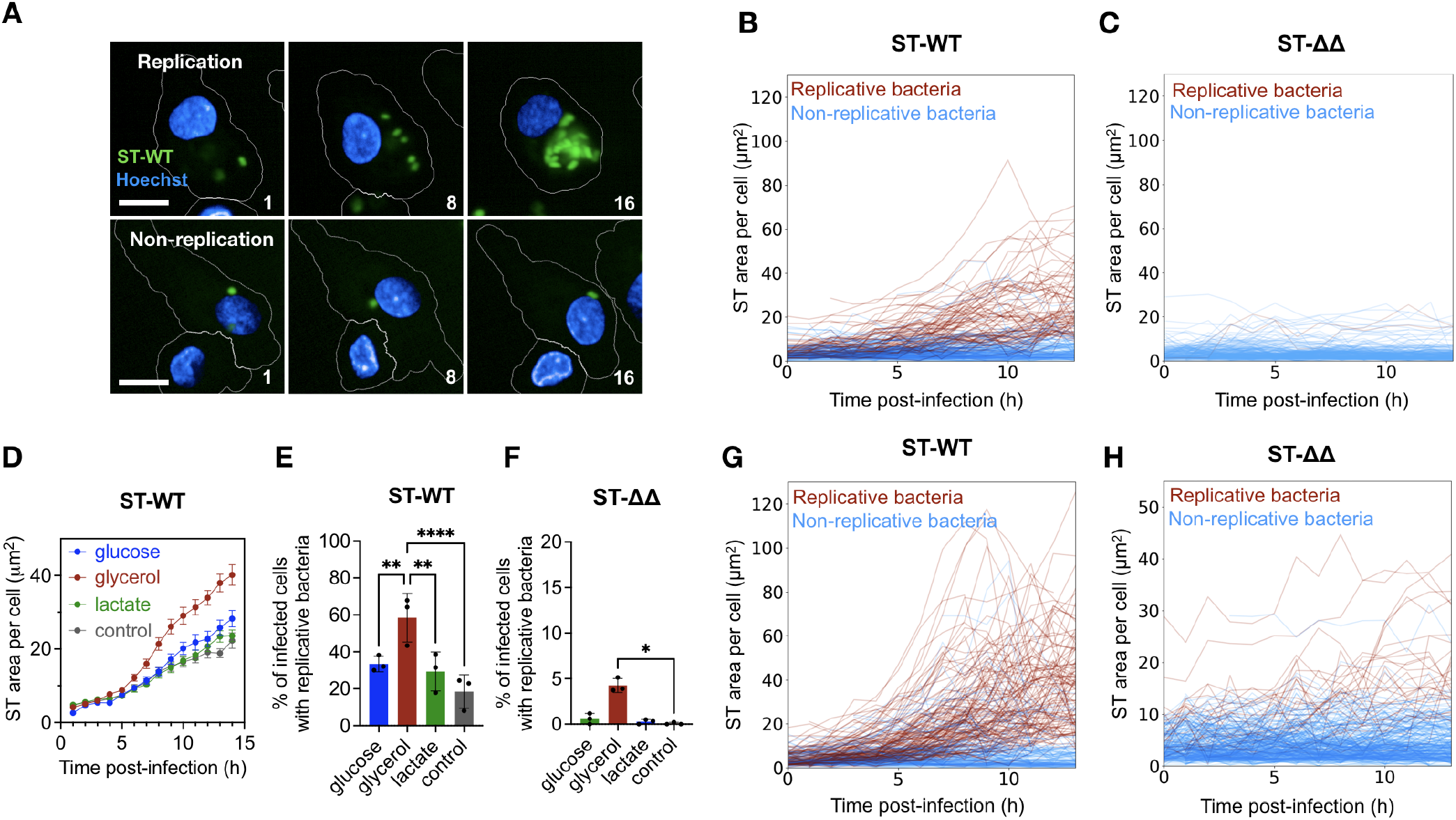
Intracellular growth dynamics of *Salmonella* Typhimurium (ST) within human monocyte-derived macrophages (hMDMs) cultured in different carbon sources. (**A**) Representative confocal images of ST-WT-infected hMDMs containing replicative or non-replicative bacteria at 1, 8 and 16 hours post-infection (hpi). GFP-expressing ST-WT bacteria are shown in green and Hoechst-stained hMDM nuclei are shown in blue. White lines delineate the cell perimeter used for segmentation. Scale bar: 10 μm. (**B**) Single-cell tracking of bacterial intracellular area (μm^2^) in ST-WT-infected hMDMs over the course of infection (1 to 16 hpi). Each line represents a single infected hMDM. Single-cell trajectories classified as hMDMs with replicative ST are shown in red, while those with non-replicative ST are shown in blue. Replication was considered if (bacterial area at t=16 hpi) (bacterial area at t=1 hpi) > 6 μm^2^. 236 single cell trajectories shown from a representative experiment. (**C**) Same as (B) but in hMDMs infected with T3SS-deficient ST-_Δ*prgH*/Δ_*ssaV* strain (ST-_ΔΔ_). 187 single cell trajectories shown from a representative experiment. (**D**) Intracellular growth dynamics of ST-WT, measured as bacterial area (μm^2^) per cell over time, in hMDMs with replicative bacteria under indicated nutrient supplementations. Data represent mean ± SEM of single cell trajectories analyzed in one experiment, representative of three independent experiments from three different donors. (**E**) Percentage of ST-WT-infected hMDMs with replicative bacteria when cultured in nutrient-limited medium supplemented with 10 mM of different carbon sources. ST-WT-infected hMDMs cultured in non-supplemented media served as control. Bars represent the mean ± SD of three independent experiments from three different donors (* = p<0.05, ** = p<0.01, ****=p<0.0001, ordinary one-way ANOVA). (**F**) Same analysis as in (E) but in hMDMs infected with ST-_ΔΔ_ strain. * = p<0.05, ordinary one-way ANOVA. (**G**) Single-cell tracking of ST-WT intracellular area (μm^2^) in infected hMDMs cultured with glycerol, with cell classification based on bacterial replication, as in (B). 261 single cell trajectories shown from a representative experiment. (**H**) Same analysis and conditions as in (G) but in hMDMs infected with ST-_ΔΔ_ strain. 253 single cell trajectories shown from a representative experiment. Please note the difference in the scale of the Y axe compared to G.

### Highly energetic macrophages support intracellular replication of *Salmonella* Typhimurium lacking both T3SSs

Surprised by the replication of the ST-Δ*prgH*/Δ*ssaV* mutant in primary macrophages cultured with glycerol, we performed similar experiments in RAW 264.7 cells, a murine macrophage cell line widely used in ST research. Due to their tumor-induced proliferative properties for their continuous cellular division, we expected that RAW macrophages should be more energetic than human primary macrophages and might support higher bacterial replication. Extracellular flux analyses using the Seahorse technology confirmed that RAW macrophages exhibited higher glycolytic and oxygen consumption rates than hMDMs **(Figure 2A)**. As expected, ST-WT showed a robust replication in RAW macrophages cultured in glucose-containing medium, with replicative bacteria growing faster than in hMDMs **(Figure 2B** and **2C)**, while the percentage of macrophages with replicative ST was similar to hMDMs **(Figure 2D)**. The ST-Δ*prgH*/Δ*ssaV* mutant replicated in 10% of infected RAW cells **(Figure 2D)**. Since T3SS-independent intracellular growth in RAW macrophages was unexpected, we measured replication in hMDMs and RAW cells by counting colony-forming units (CFUs) from infected cells, which is the traditional method used to measure bacterial replication in macrophages **(Figure S1B)**. Our results confirmed previous findings using this method ^26^ and suggested that the striking phenotype of the double T3SS-defective mutant being able to replicate intracellularly might have been overlooked. Because the percentage of cells in which the double mutant replicated was low, using CFU counting to measure bacterial replication in RAW cells might have masked this phenotype in bulk analyses **(Figure S1B)**. Live-cell tracking of bacterial replication at the single-cell level now revealed the replication phenotype of the ST-Δ*prgH*/Δ*ssaV* mutant. Our single-cell analyses showed that the double mutant generally grew at a slower rate than ST-WT, highlighting the advantage of a functional T3SS2 for intracellular replication **(Figures S1C)**. However, some ST-Δ*prgH*/Δ*ssaV* mutant bacteria reached intracellular replication levels comparable to the WT strain, showing growth of bacterial area to 50 μm^2^, which was comparable to the average growth of WT strain **(Figures 2E** and **2F)**. This suggests that the double mutant can undergo exponential intracellular growth even in the absence of T3SS-dependent secretion of bacterial effectors.

**Figure 2.**
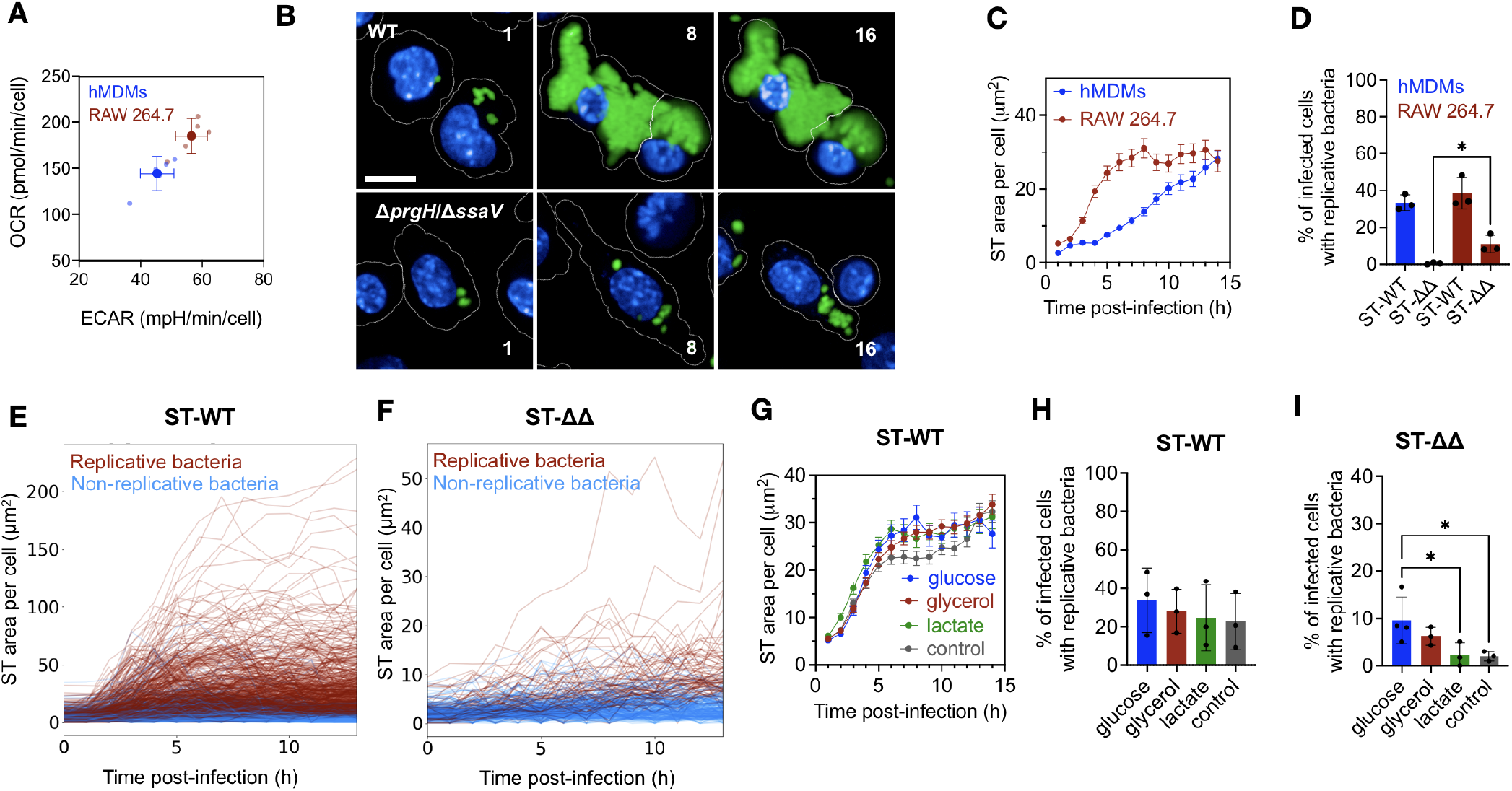
T3SS-dependent and -independent growth of ST are enhanced in high energetic RAW macrophages. (**A**) Basal extracellular acidification rate (ECAR) and basal oxygen consumption rate (OCR) of hMDMs and RAW 264.7 macrophages. Error bars indicate mean ± SD of five replicates (small circles) representative of at least three biological replicates from three different donors. (**B**) Representative confocal images of RAW macrophages infected with either ST-WT or ST-_ΔprgH/Δ_ssaV strains at 1, 8 and 16 hpi. GFP-expressing ST is shown in green and Hoechst-stained nuclei of RAW cells are shown in blue. White lines delimit cell perimeter used for segmentation. Scale bar: 10 μm. (**C**) Intracellular growth dynamics of ST-WT, measured as bacterial area (μm^2^) per cell over time, in those infected macrophages supporting Salmonella replication. (**D**) Percentage of hMDMs and RAW macrophages supporting Salmonella replication when infected with either ST-WT or ST-_ΔprgH/Δ_ssaV strain (ST-_ΔΔ_) and cultured in a glucose-supplemented medium (10 mM). The percentage was determined by single cell analysis of the intracellular bacterial area, as shown in Figure 1B. * = p<0.05, ordinary one-way ANOVA. (**E**) Single-cell trajectories of bacterial intracellular area (μm^2^) in RAW macrophages infected with ST-WT over the infection course (1-16 hpi). Each line represents a single infected cell. Trajectories of cells classified as supporting Salmonella replication are represented in red, while those classified as containing non-replicative ST are shown in blue. Single-cell analysis was performed as in Figure 1B. 957 single cell trajectories shown from a representative experiment. (**F**) Same as (E) but in RAW macrophages infected with the ST-_ΔΔ_ strain. 253 single cell trajectories shown from a representative experiment. (**G**) Effect of nutrient supplementation on the intracellular growth dynamics (μm^2^) of ST-WT in RAW macrophages showing replicative bacteria. (**H**) Effect of nutrient supplementation on the percentage of RAW macrophages supporting ST-WT replication. (**I**) Same as (H) but in RAW cells infected with ST-_ΔΔ_ strain. Data in (D), (H), and (I) represent mean ± SD of three independent experiments. Data in (C) and (G) represent mean ± SEM of single cell trajectories analyzed in one experiment, representative of three independent experiments. * = p<0.05, ordinary one-way ANOVA.

Given the relevance of key nutrients for bacterial replication in hMDMs, we examined whether these nutrients also influenced ST-infection dynamics in RAW macrophages. Glucose, glycerol and lactate slightly increased ST-WT replication in permissive RAW cells compared to control conditions **(Figure 2G)**. Overall, the percentage of cells harboring replicative bacterial vacuoles was remarkably similar across all culture conditions **(Figure 2H)**, suggesting that ST-WT replication rate in RAW cells is already near its maximal capacity independently of the supplement used. Consequently, altering host metabolism by supplementing specific carbon sources had a lower impact on bacterial replication in RAW cells compared to hMDMs **(Figure 1D** and **1E)**. In contrast, carbon source choice had a crucial effect in the intracellular growth of the ST-Δ*prgH*/Δ*ssaV* mutant strain **(Figure 2I)**. Glucose or glycerol markedly increased the percentage of cells harboring replicative ST-ΔprgH/ΔssaV bacteria compared to control conditions (10 ± 4% in glucose vs. 2 ± 1% in control, p<0.05) **(Figure 2I)**, with bacteria replicating at similar rate in both conditions **(Figure S1D)**. This contrasted with results obtained in hMDMs, where glucose did not promote the growth of the double mutant **(Figure 1F)** but glycerol did **(Figure 1H)**. In summary, our results showed that high energetic RAW macrophages are more permissive than hMDMs for the replication of the T3SS-defective strain, particularly when specific nutrients, such as glucose or glycerol, were provided to host cells.

### *Salmonella* Typhimurium lacking both T3SSs grow inside vacuoles fueled by glucose and glycerol

To explain the T3SS-independent replication of ST in RAW macrophages, we hypothesized that bacteria might escape from the SCV into a more nutrient-rich environment such as the cytoplasm of infected cells, resembling the cytoplasmic escape observed for ST-WT in epithelial cells. ^27,28^ In this scenario, the T3SS-defective ST-Δ*prgH*/Δ*ssaV* mutant might replicate intracellularly despite the absence of T3SS effectors, relying solely on nutrients available in the cytoplasm, similarly to its ability to grow at WT rates in liquid broth. To test this hypothesis, we assessed SCV integrity in RAW cells using the Galectin3mOrange (Gal3) reporter, which recognizes and binds to host glycans exposed on ruptured SCVs during infection. ^29^ Thus, by measuring the recruitment of Gal3 to the bacterial compartment, we analyzed SCV membrane rupture **(Figure 3A)**. We used HeLa cells as control and, as previously reported, ^30^ approximately 20% of ST-WT-infected HeLa cells exhibited SCV rupture **(Figure 3B)**. We observed a similar frequency of SCV ruptures in ST-WT-infected RAW macrophages at early time points post-infection **(Figure 3C)**. However, SCV rupture events were undetectable in RAW cells infected with the ST-Δ*prgH*/Δ*ssaV* mutant **(Figure 3C)**, suggesting that while ST-WT can escape the SCV in macrophages, the T3SS-defective strain remains confined within the vacuole. To confirm this, we introduced the SINA reporter into the ST-WT and -Δ*prgH*/Δ*ssaV* strains, ^31^ which enabled the localization of single bacteria within the SCV or in the cytoplasm of infected cells **(Figure 3D)**. Using this system, we classified infected cells based on ST localization **(Figures 3E, 3F** and **3G)**. As control, we used ST-WT infection of HeLa cells, where approximately 20% of infected cells harbor cytoplasmic ST-WT due to SCV escape and subsequent hyperreplication **(Figure 3E** and ref. ^31^). When RAW cells were infected with ST-WT, we observed a similar percentage of cells containing cytoplasmic bacteria as in HeLa cells **(Figure 3F)**. However, the appearance of cytoplasmic ST-ΔprgH/ΔssaV in infected macrophages was marginal, suggesting that the double mutant replicated within the SCV **(Figure 3G)**.

**Figure 3.**
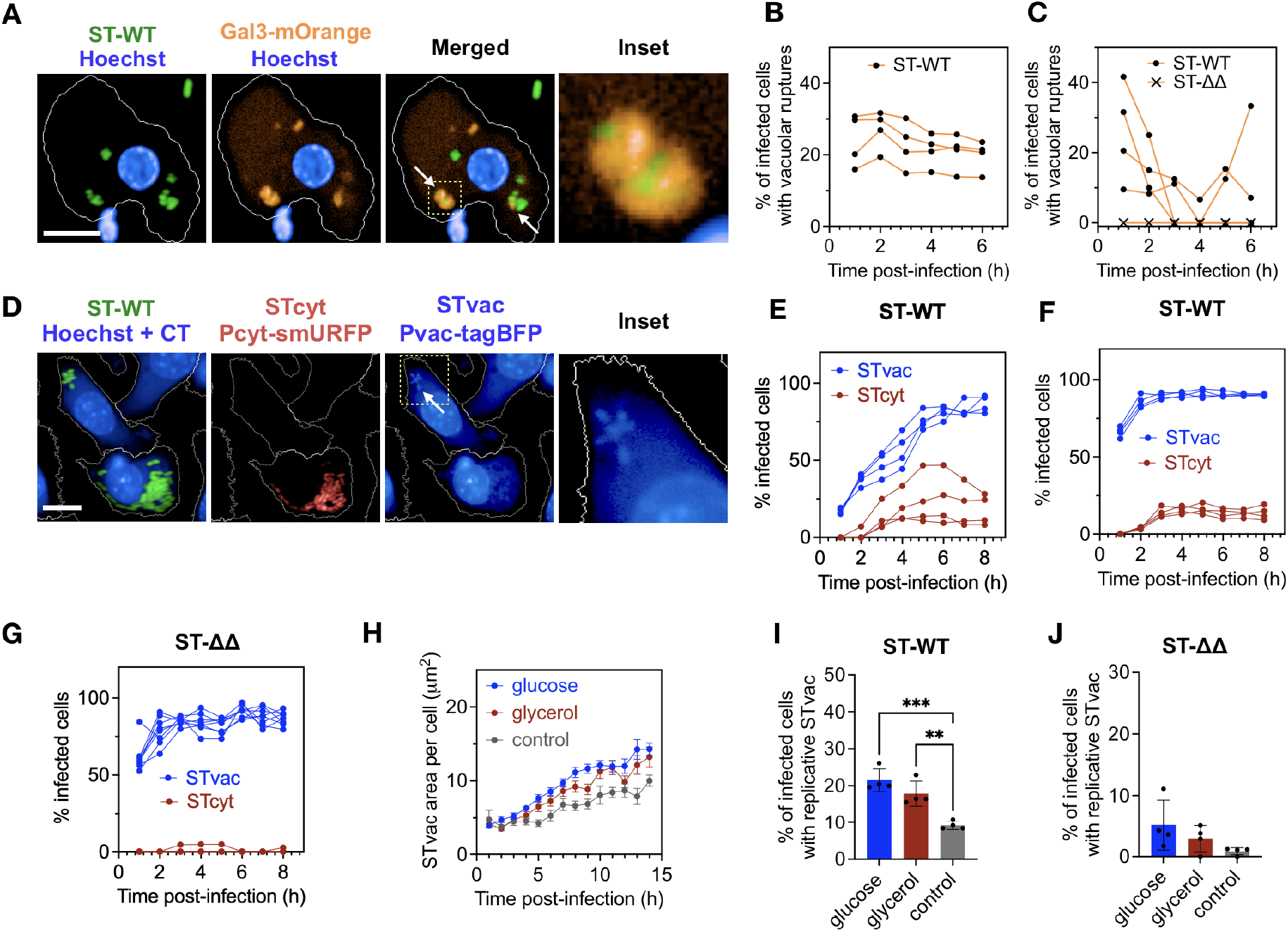
T3SS-independent ST growth occurs within SCVs. (**A**) Representative confocal images of RAW macrophages expressing Galectin-3 (Gal3)-mOrange, infected with ST. Gal3 (orange) recruitment to ST (green), indicates vacuolar ruptures (white arrows). Inset shows higher magnification of pictures showing vacuolar ruptures. Hoechst staining of nuclei is shown in blue. Scale bar: 10 μm. (**B**) Quantification of the percentage of ST-WT-infected HeLa cells displaying vacuolar ruptures over time. (**C**) Same as (B) in RAW 264.7 cells infected either with ST-WT or ST-_Δ*prgH*/Δ_*ssaV* strain (ST-_ΔΔ_). Data in (B) and (C) represent four independent replicates from a single experiment, representative of three biological replicates. (**D**) Representative confocal images of RAW macrophages infected with ST-WT expressing SINA reporter at 6 hpi. Constitutive GFP expression shows intracellular bacteria in green. Cytoplasmic *Salmonella* (STcyt) expressing Pcyt-smURFP reporter is shown in red. Vacuolar *Salmonella* (STvac) expressing Pvac-tagBFP is shown in blue (arrow). Inset highlights STvac at higher magnification. Hoechst and cell tracker blue (CT) stains host cell nuclei and the cytoplasm in blue, respectively. White lines outline the segmented cell boundaries. Scale bar: 10 μm. (**E**) Percentage of HeLa-infected cells containing STvac or STcyt bacterial populations over time. (**F**) Same as (E) in RAW macrophages infected with ST-WT. (**G**) Same as (F) in RAW cells infected with the ST-_Δ*prgH*/Δ_*ssaV* strain (ST-_ΔΔ_). Data in (E), (F) and (G) represent four independent replicates from a single experiment, representative of three biological replicates. (**H**) Impact of nutrient supplementation on intracellular replication of *Salmonella* subpopulations in RAW macrophages growing in media supplemented with glucose, glycerol or without supplementation (control). Cells were infected with ST-WT expressing SINA and the area occupied by STvac per cell (μm^2^) was measured over time. (**I**) Impact of nutrient supplementation on the percentage of ST-WT-infected RAW macrophages showing replicative STvac. (**J**) Same as (I) in ST-_ΔΔ_-infected RAW macrophages. Data in (I) and (J) represent mean ± SD of four replicates, representative of three biological replicates.

Next, we examined how nutrient availability influences ST replication and localization. We infected RAW macrophages with ST strains carrying SINA reporter and quantified bacterial growth per cell, distinguishing between cytoplasmic and vacuolar replication. Supplementation with glycerol or glucose enhanced cytoplasmic and vacuolar growth of ST-WT compared to control conditions **(Figures 3H** and **3I)**, without altering the relative distribution of bacteria between these two compartments **(Figure S2A** and **S2B)**. Glycerol or glucose also increased vacuolar growth of the ST-Δ*prgH*/Δ*ssaV* in macrophages **(Figure 3J)**. All together, our results indicated that the T3SS-deficient mutant resides within the SCV, it does not escape to the cytoplasm, and nutrient availability plays a critical role fueling its growth within the SCV.

### Nutrient availability, rather than the glycolytic activity of host cells, drives T3SS-independent *Salmonella* replication in macrophages

Previous studies have reported that glycolysis is essential for efficient ST replication within the SCV. ^13,15,22,32^ Our results showed that glucose significantly enhanced ST replication in hMDMS and RAW macrophages **(Figure 1** and **2)** and, compared to human primary macrophages, the higher glycolytic phenotype of RAW cells correlated with a higher replication of ST **(Figure 2C** and **2D)**. Thus, we investigated whether macrophages growing in glycerol **(Figure 1D** and **1E)** promoted intracellular bacterial growth by enhancing the glycolytic pathway. We measured glycolysis in macrophages under different nutrient conditions using the Seahorse technology using 2-deoxy-D-glucose (2-DG), a well known inhibitor of glycolysis, as a control. Our results showed that glycerol indeed decreased host glycolysis **(Figure 4A)**. As glycerol supported ST growth within the SCV at maximum levels **(Figure 3H)**, we hypothesized that intracellular replication of ST was not enhanced by glycolysis itself, but rather by the overall availability of readily metabolizable nutrients within the host cell. To test this hypothesis, we measured bacterial growth under conditions where glycolysis was inhibited by 2-DG. At high 2-DG concentration (50 mM), which showed the highest glycolysis inhibition **(Figure 4A)**, the percentage of macrophages with replicative *Salmonella* and bacterial growth were decreased **(Figure 4B** and **4C)**. Low 2-DG concentration (1 mM) only partially inhibited glycolysis **(Figure 4A)** and had no effect on ST replication **(Figure S2C)**. When we provided glycerol to macrophages treated with the high concentration of 2-DG, ST replication was restored to levels comparable to those observed in glucose-fed cells **(Figure 4B** and **4C)**, strongly suggesting that glycolysis was not essential for ST growth in the SCV as long as an alternative carbon source to glucose, such as glycerol, was available.

**Figure 4.**
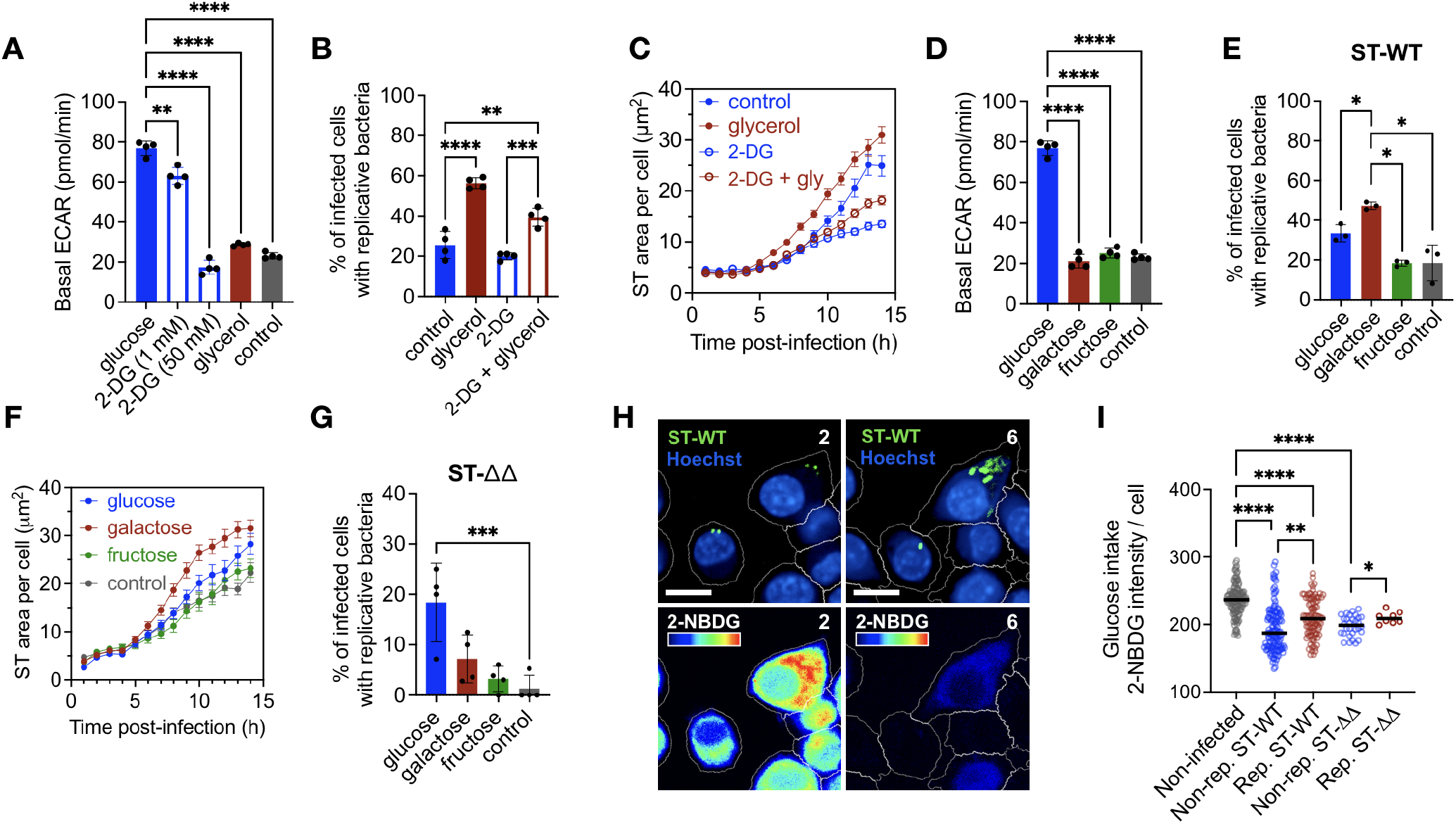
Nutrient availability, rather than the glycolytic activity of host cells, drives T3SS-dependent and -independent ST replication in macrophages. (**A**) Basal extracellular acidification rate (ECAR) of RAW macrophages cultured in 10 mM glucose, 10 mM glycerol, or 10 mM glucose supplemented with 1 mM or 50 mM of 2-DG to inhibit glycolysis. Cells in non-supplemented media (with no glucose), served as a control for low glycolysis rate. Data represent mean ± SD of five replicates, representative of at least three independent experiments. (**B**) Percentage of ST-WT-infected hMDMs supporting bacterial replication when cultured in glucose (10 mM) with or without 2-DG (50 mM), in glycerol (10 mM), or in 2-DG and glycerol. Data represent mean ± SD of three biological replicates. (**C**) Intracellular ST-WT growth dynamics measured as bacterial area (μm^2^) per cell over time in hMDMs supporting bacterial replication under the indicated nutrient conditions. Data show mean ± SEM of three independent experiments. (**D**) Basal ECAR of RAW macrophages cultured in 10 mM of glucose, galactose or fructose. Non-supplemented media was used as control. (**E**) Percentage of ST-WT infected hMDMs supporting Salmonella replication under different sugar supplementation conditions. (**F**) Intracellular ST-WT growth dynamics in infected hMDMs supporting bacterial replication, under the indicated sugar supplementation conditions. (**G**) Same as (E) in RAW macrophages infected with ST-_Δ*prgH*/Δ_*ssaV* strain (ST-_ΔΔ_). (**H**) Representative confocal images of RAW macrophages incubated for 4 h with 2-NBDG to measure glucose uptake, and infected with ST-WT. Hoechst staining of nuclei is shown in blue, bacteria in green and fluorescent glucose as a cold-hot palette. Same cells are shown at 2 hpi and 6 hpi. Scale bar: 10 μm. (**I**) Single-cell analysis of glucose uptake, measured by 2-NBDG fluorescence intensity (AU), in non-infected, ST-WTand ST-_ΔΔ_-infected macrophages at 2-hpi. Infected macrophages were classified as cells later harbouring replicative or non-replicative ST-WT or ST-_ΔΔ_. Bars represent the mean of one experiment, representative of three biological replicates. Statistical significance is indicated as *p < 0.05, **p < 0.01, ***p < 0.001, ****p < 0.0001 by ordinary one-way ANOVA.

When sugars, such as galactose or fructose, are provided as alternative carbon sources to glucose, they can enter into the glycolytic pathway at different levels. This create metabolic bottlenecks that lead to reduced glycolytic rate and the accumulation of the corresponding sugar in the cell. ^33^ Thus, to investigate whether the availability of alternative sugars can fuel bacterial growth independently of glycolysis, we supplemented macrophages with galactose or fructose. As expected, supplementing macrophages with any of these two sugars reduced glycolysis **(Figure 4D)**. Interestingly, even if both sugars reduced glycolysis, ST replication rates were very different under these two conditions. While ST replication was reduced in fructose-supplemented macrophages, bacterial growth was significantly enhanced in macrophages cultured with galactose **(Figure 4E** and **4F)**. Moreover, T3SS-independent ST replication was also observed in macrophages cultured with galactose **(Figure 4G)**. These findings reinforced our hypothesis that the accumulation of specific nutrients in the infected cell, rather than glycolytic activity, drives ST replication. To directly assess this in single cells, we used 2-NBDG and 2-FBDG to track glucose and fructose uptake, respectively, as they are fluorescent analogs of these sugars that accumulate in the cytoplasm of cells upon uptake. We measured sugar uptake at the onset of infection and then analyzed bacterial replication dynamics for each individual infected macrophage. Using this approach we classified each infected macrophage according to intracellular bacterial replication and then correlated ST replication with host cell sugar intake at the single-cell level **(Figure 4H** and **4I)**. Our analyses showed that, at times prior to bacterial replication, macrophages displaying a higher glucose uptake were those that supported bacterial growth at later time points, in contrast to non-permissive macrophages, which exhibited a drop in glucose intake **(Figure 4H)**. Consistent with fructose being unable to enhance ST replication **(Figure 4E)**, no differences were found in fructose uptake between macrophages supporting bacterial replication and those macrophages where ST did not replicate **(Figure S2D)**, suggesting that, unlike glucose, fructose uptake and availability at the infected cell did not promote ST replication in macrophages. All together, our results showed that intracellular ST replication in macrophages was primarily dictated by the presence of specific nutrients, rather than the glycolytic activity of host cells. Thus, within the heterogeneity of macrophages infected by ST, only those with higher accumulation of preferred ST nutrients, such as glucose, glycerol or galactose, promoted the intracellular replication of ST, which can occur independently of host glycolysis and the action of T3SS bacterial effectors.

## Discussion

In this study we demonstrated that providing macrophages with preferred nutrients allows ST to replicate intracellularly without the need for functional T3SSs (T3SS1 and T3SS2). T3SS-dependent effectors are considered essential for *Salmonella* intracellular survival, allowing bacteria to remodel host phagosomes into specialized replication niches. ^5,8–12^ Surprisingly, our findings challenge this paradigm by showing that under nutrient-rich conditions, *Salmonella* mutants deficient in both T3SS1 and T3SS2 (ST-Δ*prgH*/Δ*ssaV* mutant strain) can replicate within macrophages, indicating that access to nutrients alone can partially substitute for T3SS-mediated virulence.

These observations align closely with recent *in vivo* studies emphasizing that the pathogenic strategy of ST relies on its metabolic versatility and ability to exploit diverse host nutrients. ^13,15,34,35^ Steeb and colleagues previously highlighted that *Salmonella* virulence *in vivo* depends on the parallel utilization of multiple carbon sources including glucose, glycerol, lactate, fatty acids, gluconate, arginine, and N-acetylglucosamine. ^13^ Importantly, glucose was considered uniquely crucial for systemic infections, yet the collective contribution of these nutrients appears necessary to support optimal pathogen growth in host tissues. Our results extend this understanding by demonstrating at the single-cell level that the macrophage nutrient status critically determines whether intracellular *Salmonella* replicate, independently of T3SS functions. We showed that providing macrophages with glycerol, one of ST preferred carbon sources, bypasses the necessity of the bacterium of increasing host cell glycolysis and the actions of its T3SSs effectors. This supports a model where nutrient availability is not simply a downstream consequence of T3SS actions but can independently define bacterial growth within host cells.

The vacuolar compartment *Salmonella* inhabits is typically nutrient-limited, requiring T3SS effectors to create an environment suitable for bacterial proliferation. ^36^ The T3SS2 from ST facilitates nutrient acquisition by remodeling the SCV and establishing direct nutritional links with host endosomal compartments through the *Salmonella*-induced filaments (SIFs). ^36^ By providing abundant nutrients externally, this seems to effectively bypass the need for remodeling. This suggests that one of the primary purposes of T3SS effector proteins is to overcome host-imposed nutrient restriction within macrophage vacuoles. Recent research underscores this mechanism, showing that SPI-2 mutants exhibit metabolic stress inside macrophages due to their inability to access essential metabolites sequestered in endolysosomal compartments. ^36^ Beyond access to nutrients, *Salmonella* also actively reprograms host cell metabolism using T3SS effectors to create a favorable intracellular niche. Recent studies have shown that *Salmonella* effectors manipulate macrophage metabolic pathways to boost nutrient availability. ST infection drives murine macrophages toward a Warburg-like aerobic glycolysis phenotype, characterized by high glucose uptake and lactate production. ^22,37^ The T3SS1 effector SopE2 was shown to play a key role in this metabolic reprogramming by blocking the host serine biosynthesis pathway, which in turn causes an accumulation of upstream glycolytic intermediates such as 3-phosphoglycerate (3PG), pyruvate, and lactate. ST can directly exploit these changes by consuming 3PG and lactate as intracellular nutrients. ^22,37^ Indeed, lactate accumulation synergized with the T3SS2 effector SteE to drive macrophages towards an M2like, anti-inflammatory polarization, and this shift was required for the lactate-mediated boost in bacterial growth. ^37^ Our results in human macrophages showed that supplementing macrophages with lactate or pyruvate had no effect on ST intracellular growth. Indeed, one key insight from our study is that the extent to which nutrient supplementation rescued intracellular replication varied markedly between macrophage types. While RAW macrophages showed robust support for T3SS-deficient ST replication, human primary macrophages were more restrictive. Human macrophages supported some replication of the T3SS-deficient ST strain under nutrient-rich conditions, such as glycerol, but not to the same levels as RAW cells. This suggests that the capacity of the host cell to take up and share nutrients with the pathogen is a key determinant of the efficient replication of ST. Moreover, Rosenberg and colleagues demonstrated that *Salmonella* can sense and exploit host-derived metabolites such as succinate, a metabolite that accumulates in macrophages upon activation ^38^. Host-derived succinate does not only serve as a nutrient but also acts as a crucial activation signal, promoting antimicrobial resistance and the expression of key virulence factors, which illustrates the complexity of the metabolic interplay. In our study, the elevated glycolytic flux of RAW macrophages likely resulted in enhanced uptake and vacuolar delivery of extracellular metabolites, such as glucose, galactose or glycerol, significantly favoring intracellular bacterial growth. Overall, our results support a model where the nutritional state of the macrophage can override the requirement for T3SS, highlighting the role of host metabolism in controlling intracellular infections.

The main strength of our study is, ironically, its main limitation. By using single-cell tracking of bacterial dynamics in infected macrophages, our approach was uniquely able to show the surprising replication phenotype of the T33Sdeficient ST strain. Thus, the main limitation of our study is the obligatory *in vitro* settings needed to track infection dynamics in individual macrophages. Future research should include *in vivo* validation to confirm whether nutrient abundance similarly influences T3SS dependency within infected tissues. Employing diabetic animal models or *in vivo* metabolic modulation approaches would be useful for assessing how physiological changes in host metabolism impact *Salmonella* pathogenicity. Additionally, a deeper mechanistic understanding of how extracellular or cytosolic metabolites reach intravacuolar bacteria remains essential. Advanced imaging coupled to metabolic tracing methods, which ideally would render single-cell analyses of infected cells, could clarify whether nutrients enter via vacuolar membrane disruption, active host transport, or passive diffusion.

In summary, our work revealed nutrient availability as a critical determinant for circumventing the necessity of T3SS-mediated virulence of *Salmonella*. By highlighting hostpathogen metabolic interactions as central to intracellular survival, these findings inspire the development of innovative metabolic-targeted therapies limiting pathogen growth by restricting intracellular nutrient accessibility.

## Acknowledgements

We acknowledge Carmen Buchrieser for the critical reading of the manuscript and her support, and to all C.B’s lab members for fruitful discussions. We thank Nathalie Aulner, Anne Danckaert, Nassim Mahtal and the Photonic BioImaging (PBI) UTechS at Institut Pasteur for their support. We are grateful to the healthy volunteers for their participation in the study. We acknowledge Hélène Laude, the ICAReB-Clin of the Medical Direction and the ICAReB-Biobank of the Biological Resource Center of the Institut Pasteur for providing blood samples from healthy volunteers, managing the participants’ visits, and preparing the blood samples from donors. This research was funded by the Institut Pasteur, the Agence National de Recherche (ANR21-CE15-0038-01 to P.E.), the Programmes Transversaux de Recherche (PTR-651) from Institut Pasteur to P.E and the Région Ile-de-France (program DIM1Health) to PBI (part of FranceBioImaging, ANR-10INSB-04–01), and the MCIN/AEI/10.13039/501100011033/ from Spanish government (grant PID2022-136863NB-I00) to J.B.B.

## Methods

### Bacterial strains and plasmids

All *S*. Typhimurium strains used in this study are derived from the parental strain ATCC 14028 and are listed in Table 1. Bacteria were routinely cultured in Lysogeny broth (LB) supplemented with ampicillin 100 μg/mL. For pFPV25-mNeptune plasmid, mNeptune synthetic gene was amplified by PCR using 5’ and 3’ oligos flanked by XbaI and HindIII sites, respectively. XbaI/HindIII digested PCR product was cloned into XbaI/HindIII digested pFPV25.1 vector at the place of GFP. Construct was verified by sequencing. ST-Δ*prgH*/Δ*ssaV* derivative mutant was verified by PCR amplification using specific *ssaV* and *prgH* external primers: ssaV5’=CGAGCTCTGGTTACGATTAC, ssaV3’=CAGCCTCAGACGTTGCATC, PprgHfwEco=AGTCGAATTCTCGTGATTATTGCTAATCG, PprgHrvEco=AGTCGAATTCATATACTGTTAGCGATGTC. Amplification of the external regions were performed on a T100 Thermal Cycler (Bio-Rad) using MyTaq Red DNA polymerase (Bioline) according to the fabricant instructions. 14028 wild type and steD strains were used as controls of the expected size of *prgH* and *ssaV* genes.

**Table 1.** List of bacterial strains and plasmids used in this study. * Channel (excitation/emmision, in nm): Blue = 375/435-480, Green = 488/500-550, Red = 561/570-630, Far Red = 640/650-760.

| Strain/Plasmid | Relevant genotype/description | Antibiotic Selection | Channel* | Ref. |
| --- | --- | --- | --- | --- |
| ST-WT-GFP (SV9760/pFPV25.1) | Wild-type *Salmonella enterica* ser. Typhimurium 14028 | Ampicillin | Green | 39 |
| ST-$\Delta prgH\Delta ssaV$-GFP (SV6747/pFPV25.1) | T3SS1- and T3SS2-defective ST 14028 | Ampicillin | Green | This study |
| ST-$\Delta prgH$-GFP (SV5379/pFPV25.1) | T3SS1-defective ST 14028 | Ampicillin | Green | 39 |
| ST-$\Delta ssaV$-GFP (SV5136/pFPV25.1) | T3SS2-defective ST 14028 | Ampicillin | Green | 39 |
| ST-WT-Neptune | Wild-type *Salmonella enterica* ser. Typhimurium 14028 | Ampicillin | Far-Red | This study |
| ST-$\Delta prgH\Delta ssaV$-Neptune | T3SS1- and T3SS2-defective ST 14028 | Ampicillin | Far-Red | This study |
| ST-WT-SINA | Wild-type *Salmonella enterica* ser. Typhimurium 14028 | Ampicillin | Blue, Green, Red, Far-Red | This study |
| ST-$\Delta prgH\Delta ssaV$-SINA | T3SS1- and T3SS2-defective ST 14028 | Ampicillin | Blue, Green, Red, Far-Red | This study |
| pFPV25.1 | GFP, constitutive | Ampicillin | Green | 40 |
| pFPV25-mNeptune | pFGV25, mNeptune, constitutive | Ampicillin | Far-Red | This study |
| pSINA1.1. | SINA 1.1. reporter, Pconst-Timer, Pvac-tagBFP, P-smURFP | Ampicillin | Blue, Green, Red, Far-Red | 31 |
| pmOrange-Galectin3 | mOrange-Galectin3, constitutive expression in eukaryotic cells | Kanamycin | Red | 41 |

#### Host cells

Human monocyte-derived macrophage (hMDMs) isolation and differentiation was performed as previously described. ^42^ Human peripheral blood samples were collected from healthy volunteers through the ICAReB-Clin (Clinical Investigation platform) of the Institut Pasteur. All participants received an oral and written information about the research and gave written informed consent in the frame of the healthy volunteers CoSImmGEn cohort (Clinical trials NCT 03925272), after approval of the CPP Ile-de-France I Ethics Committee (2011, January 18th). Peripheral blood mononuclear cells (PBMCs) were isolated by Ficoll density gradient centrifugation (Lymphocyte Separation Medium, Eurobio Scientific) at room temperature. Monocytes were positively sorted using CD14 magnetic beads (CD14 MicroBeads, Miltenyi Biotec). Following, monocytes were differentiated into hMDMs by culturing them in X-VIVO15 medium without Phenol Red (Lonza) in Nunc Up-Cell plates (Thermo Scientifics) and in the presence of recombinant human M-CSF (rh-MCSF, Miltenyi Biotec) at a concentration of 25 ng/ml for 6 days at 37 °C with 5% CO_2_ in a humidified incubator. The day before infection, cells were detached from UpCell plates and plated in 384-well plates (Greiner Bio-One) at a cell density of 10,000 cells per well in X-VIVO15. The RAW 264.7 macrophage-like cell line (ATCC TIB-71) was obtained from ATCC and cultured in RPMI medium supplemented with 10% (v/v) heat-inactivated fetal bovine-serum (FBS) (Gibco). HeLa cells (ATCC CCL-2) were obtained from ATCC and cultured in DMEM medium supplemented with 10% (v/v) heat-inactivated FBS. HeLa GAL3-mOrange-cells were a gift from Dr. Jost Enninga.

#### Infections of host cells with *S*. Typhimurium

A bacterial colony from LB agar plate supplemented with 100 μg/mL ampicillin was used to inoculate a culture in LB medium containing 100 μg/mL ampicillin and grown shaking overnight at 37 °C. On the day of the infection, 150 μL of the overnight culture was sub-cultured in 5 mL of fresh LB medium (1:33 dilution) and incubated with shaking for 3 hours, allowing the bacteria to reach an early stationary phase. Cultures were then washed in FBS-free cell culture medium. hMDMs and RAW 264.7 macrophages were infected with ST-WT at a multiplicity of infection (MOI) of 10, ST-ΔprgH/ΔssaV at an MOI=50, while HeLa cells were infected with ST-WT at an MOI=50. Infected cells were centrifuged at 200 xg for 5 min to synchronize infection, followed by 30 min incubation at 37 °C and 5% CO_2_. To remove extracellular bacteria, cells were washed five times with PBS, and fresh media containing 100 μg/ml of gentamicin was added for 20 min. Then, the medium was replaced with fresh culture media containing 10 μg/mL gentamicin, which was maintained for the duration of the infection to avoid ST extracellular replication. For the supplementation with the different carbon sources, one hour before infection, the standard cell culture medium was replaced by RPMI medium (without glucose, glutamine, or pyruvate, Agilent Biosciences), containing 10% (v/v) FBS (Gibco) and supplemented with 10 mM of each corresponding carbon source. Same media was used to cultured infected macrophages during the course of infection. Infections were performed with RPMI medium + FBS without any additional supplementation.

#### Staining of living cells and automated time-lapse confocal imaging acquisition

For automated time-lapse confocal imaging experiments, we performed infection assays in 384well microplates as previously described. ^42–44^ Each experiment was performed using four technical replicates and repeated to obtain at least three biological replicates. To track individual cells, nuclei and cytoplasm were stained one hour before infection with 200 ng/mL Hoechst H33342 (H3570, Life Technologies) and 25 μM CellTracker Blue (C12881, Invitrogen), respectively. ^25^ Cells were washed before infection to prevent bacterial staining. Image acquisition was performed using an automated microlens-enhanced spinning disc confocal microscope (Opera Phenix High Content Screening System, PerkinElmer) equipped with a 63x water objective. Excitation lasers operated at 405, 488, 561, and 640 nm, with emission filters set at 450, 540, 600, and 690 nm. To capture the dynamics of infection, images of multiple fields (ranging from 10 to 16) were acquired every hour over a 16-hour period in an incubation chamber maintained at 37 °C with 5% CO_2_. For glucose and fructose intake assays, after 2 h post-infection, 2-NBDG (Life Technologies) or 2-NBDG (Cayman Chemical) was added at a final concentration of 0.5 mM. After 30 min of incubation cells were washed 3 times in PBS and fresh media containing either glucose or fructose (10 mM) was added and maintained during the course of infection.

#### Image Analyses

Image analyses were performed using the Harmony software v.4.9 (Perkin Elmer) and self-built scripts (shared upon request). Briefly, cell nuclei was segmented using Hoechst signal, cytoplasm region using Cell Tracker Blue background, intracellular bacteria were identified using GFP/RFP/Neptuno signal (depending the assay), and metabolic parameters such as 2-NDBG or 2-FDBG sugar uptake were measured using green signal. Reconstruction of single-cell tracks and backtracking analyses were performed using BATLI software as described previously. ^25^

#### Real-time extracellular flux assays (Seahorse)

ECAR and OCR of macrophages were determined using an XF96 extracellular flux analyzer (Seahorse Bioscience, Agilent). hMDMs (6 × 10^5^ cells/well) or RAW 264.7 macrophages (4 × 10^5^ cells/well) were plated in XF96-cell culture plates containing XVIVO15 or RPMI supplemented with 10% (v/v) FBS, respectively. One hour before analysis, the culture medium was removed from the cells, and the cells were incubated at 37 °C and 0% CO_2_ in XF assay RPMI medium containing 10% (v/v) FBS and supplemented with 10 mM of each corresponding carbon source. For metabolic analyses of infected macrophages, cells were incubated one hour before infection in RPMI medium + 10% FBS supplemented with the corresponding carbon source and then infected with ST for 2 hours as described above before performing metabolic assays. ECAR and OCR values were taken before and after the sequential addition of glucose (10 mM), Oligomycin (1 μM) and 2-DG (50 mM) (XF Glycolysis Stress Test Kit, Seahorse Bioscience, Agilent). Basal ECAR and OCR values were obtained by subtracting the non-glycolytic ECAR values and non-mitochondrial OCR values, respectively, and after normalizing to the cell number.

#### Colony-forming units (CFUs) counting

hMDMs (6 × 10^5^ cells/well) or RAW 264.7 macrophages (4 × 10^5^ cells/well) were plated in 96-well plates. Infections were performed as described before. At 20 hpi, cells were lysed with 100 μl of sterile deionized water and plated in LB agar plates. After one day of incubation at 37 °C, CFUs were counted.

#### Generative AI statement

During the preparation of this work, authors used ChatGPT 4 (OpenAI) and Gemini 2.5 (Google) Large Language Models (LLMs) in order to correct grammar and flow of some parts of the text. After using these tools, authors reviewed and edited the content as needed and take full responsibility for the content of the publication.

## Supplementary Information

### Supplementary Figures

**Figure S1.**
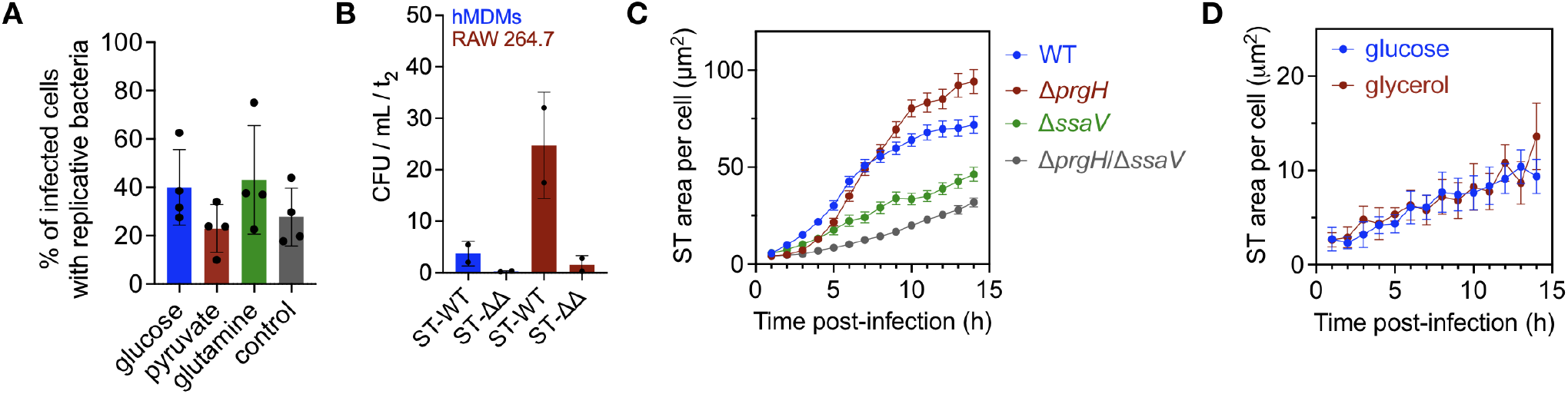
Related to Figure 1 and 2. (**A**) Percentage of ST-WT-infected hMDMs with replicative bacteria when cultured in nutrient-limited medium supplemented with 10 mM of different carbon sources. ST-WT-infected hMDMs cultured in non-supplemented media served as control. Bars represent the mean ± SD of three independent experiments from three different donors (*=p<0.05, **=p<0.01, ****=p<0.0001, ordinary one-way ANOVA). (**B**) CFU counts of hMDMs infected with ST-WT or ST-Δ_prgH/Δ_ssaV strain at 20 hpi. (**C**) Intracellular growth dynamics (μm^2^) of ST-WT, ST-_Δ_prgH, ST-_Δ_ssaV or ST-_ΔprgH/Δ_ssaV strains in RAW macrophages showing replicative bacteria. (**D**) Effect of nutrient supplementation on the intracellular growth dynamics (μm^2^) of ST-_ΔprgH/Δ_ssaV in RAW macrophages showing replicative bacteria.

**Figure S2.**
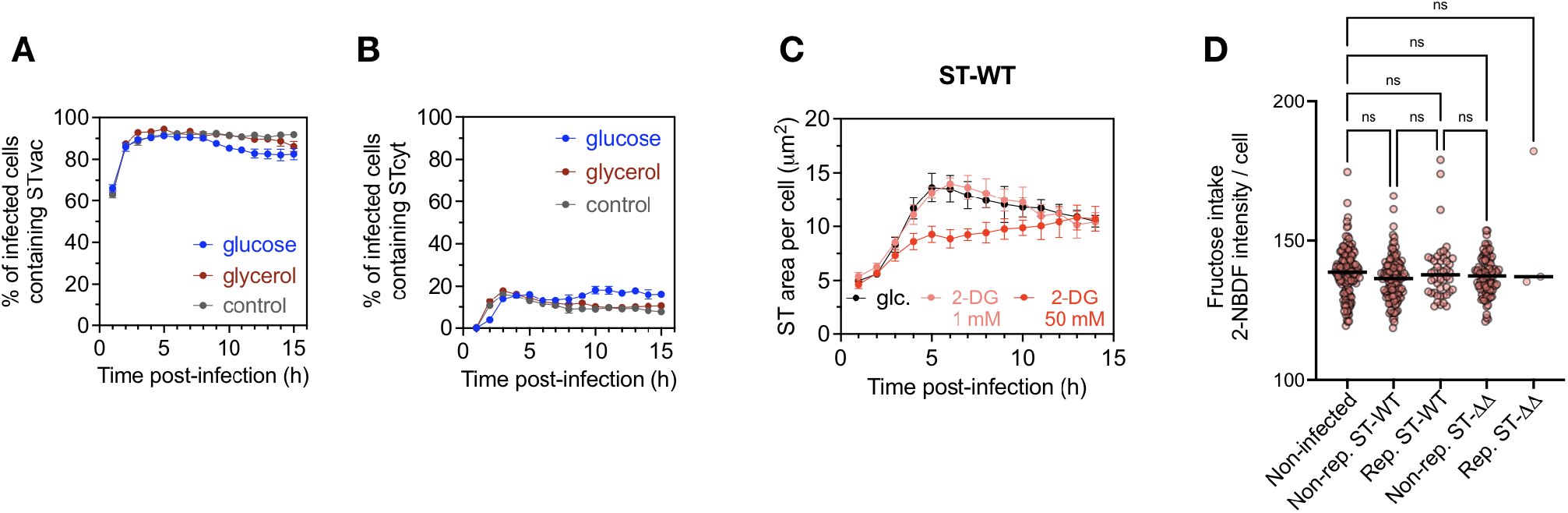
Related to Figure 3 and 4. (**A**) Percentage of infected macrophages with ST in vacuoles (STvac) under nutrient supplementation as indicated. (**B**) Same as (A) for ST in the cytoplasm (STcyt). (**C**) Intracellular ST-WT growth in RAW macrophages during 2-DG high- and low-concentration treatments, glc. indicates control conditions with glucose. (**D**) Single-cell analysis of fructose uptake, measured by 2-NBDF fluorescence intensity (AU), in non-infected, ST-WT-, and ST-_ΔΔ_-infected macrophages at 2-hpi. Infected macrophages were classified as cells later harbouring replicative or non-replicative ST-WT or ST-_ΔΔ_. Bars represent the mean of one experiment, representative of three biological replicates. Statistical significance is indicated as *=p<0.05, **=p<0.01, ***=p<0.001, ****=p<0.0001 by ordinary one-way ANOVA.

## Notes

### Competing Interest Statement

The authors have declared no competing interest.

